# Mycobacteriophage D29-Derived LysB Enhances Anti-Tubercular Therapy in Experimental Pulmonary Tuberculosis

**DOI:** 10.64898/2026.05.28.728497

**Authors:** Sunil Kumar Raman, Rahul Sharma, Rutuja Gangakhedkar, Priyamvada Nath, Amit Misra, Vikas Jain, Amit Kumar Singh

**Affiliations:** Pharmaceutics and Pharmacokinetics Division, CSIR-Central Drug Research Institute, Lucknow 226031, India; Experimental Animal Facility, ICMR-National JALMA Institute for Leprosy & Other Mycobacterial Diseases, M. Miyazaki Marg, Tajganj, Agra-282004, Uttar Pradesh, India; Microbiology and Molecular Biology Laboratory, Indian Institute of Science Education and Research, Bhopal – 462066, India; Laboratory of Antimicrobial Innovation, School of Interwoven Arts and Sciences, Krea University, Sri City – 517646, Andhra Pradesh; Assistant Professor, Academy of Scientific and Innovative Research (AcSIR), Ghaziabad, Uttar Pradesh 201002, India; Amit Kumar Singh, Scientist-E, Experimental Animal Facility Division, ICMR-National Institute for Research in Tuberculosis (NIRT-JALMA North India Campus), Agra, U.P., India

**Author notes:** Correspondence: Dr. Amit Kumar Singh, Scientist-E, Experimental Animal Facility, ICMR-National JALMA Institute for Leprosy & Other Mycobacterial Diseases, M. Miyazaki Marg, Tajganj, Agra-282004, Uttar Pradesh, India, Dr. Vikas Jain, Laboratory of Antimicrobial Innovation, School of Interwoven Arts and Sciences, Krea University, Sri City – 517646, Andhra Pradesh, India. Contributed Equally (Sunil Kumar Raman, Rahul Sharma).

**Keywords:** *Mycobacterium tuberculosis*, D29 LysB, mycolylarabinogalactan esterase, intranasal delivery, Adjunctive therapy

## Abstract

Adjunctive therapies that enhance the efficacy of existing antitubercular drugs are needed for drug-resistant tuberculosis. We evaluated the efficacy of intranasally administered recombinant D29 LysB, a mycobacteriophage-derived mycolylarabinogalactan esterase, in murine and guinea pig models of pulmonary tuberculosis. BALB/c mice and guinea pigs were aerosol-infected with *Mycobacterium tuberculosis* H37Rv and treated for 4 weeks with LysB alone or with standard antitubercular therapy (ATT: rifampicin, isoniazid, pyrazinamide). Outcomes included pulmonary and extrapulmonary bacterial burden (CFU), lung and spleen histopathology, cytokine profiling, and humoral immune responses. LysB monotherapy produced modest pulmonary CFU reductions. When given adjunctively with ATT, LysB produced an additional 0.6–0.7 log^10^ reduction in lung CFU compared with ATT alone and decreased splenic dissemination in both species. Combination therapy improved tissue pathology, reducing granulomatous involvement and preserving pulmonary architecture. LysB treatment increased TNF-α with a moderate rise in IL-10, a profile consistent with enhanced antibacterial immunity without excessive inflammatory damage. Repeated intranasal administration was well tolerated; no IgE-mediated hypersensitivity was detected. LysB-specific IgG developed but did not diminish therapeutic efficacy. These results show that intranasal D29 LysB augments the bactericidal and histopathological effects of standard ATT *in vivo* and support further development of inhaled phage-derived lysins as adjunctive therapies for drug-resistant tuberculosis.

**Importance:** Tuberculosis remains a major cause of infectious mortality worldwide, and the increasing burden of multidrug-resistant and extensively drug-resistant disease continues to challenge effective treatment. New therapeutic approaches that complement conventional antibiotics are urgently needed. In this study, intranasally delivered recombinant mycobacteriophage-derived LysB was well tolerated and enhanced treatment efficacy in experimental pulmonary tuberculosis. Adjunctive LysB improved bacterial clearance, reduced tissue pathology, and modulated host immune responses in both murine and guinea pig models. These findings highlight phage-derived endolysins as promising inhalable adjunctive therapeutics for drug-resistant tuberculosis.

## Introduction

Tuberculosis (TB), caused by *Mycobacterium tuberculosis* (*M. tuberculosis*), remains a leading cause of infectious mortality worldwide despite decades of therapeutic advances. The disease continues to impose a major public health burden, particularly in low- and middle-income countries, where incidence and mortality remain disproportionately high. According to the World Health Organization, approximately 10.7 million new TB cases were reported globally, resulting in nearly 1.3 million deaths [1]. Efforts to control TB are increasingly challenged by the emergence and spread of drug-resistant disease, particularly multidrug-resistant (MDR) and extensively drug-resistant (XDR) strains, which substantially complicate treatment outcomes. Current anti-tuberculosis regimens are prolonged, complex, and frequently associated with significant toxicity, underscoring the urgent need for novel therapeutic strategies that can complement conventional antibiotics or improve their efficacy [1, 2].

Bacteriophages and their lytic enzymes have emerged as promising alternatives for combating bacterial infections, particularly those caused by antibiotic-resistant pathogens [3, 4]. Among these, mycobacteriophages specifically infect mycobacterial species and have attracted considerable interest as potential therapeutic tools against diseases such as TB [5–7].

Mycobacterial lysis is mediated by phage-encoded endolysins, of which LysA and LysB are key functional enzymes. LysA hydrolyzes the peptidoglycan layer, whereas LysB, a mycolylarabinogalactan esterase, cleaves the ester linkage between mycolic acids and arabinogalactan within the mycobacterial cell envelope [8, 9]. Disruption of this highly specialized envelope compromises bacterial integrity and increases susceptibility to host immune defenses and antibiotic-mediated killing [8].

Emerging evidence supports the therapeutic potential of recombinant LysB derived from mycobacteriophage D29. *In vitro* studies have demonstrated potent antimycobacterial activity against *Mycobacterium abscessus*, *Mycobacterium ulcerans*, and multidrug-resistant *M. tuberculosis*, while combination treatment with conventional antibiotics has shown additive or synergistic effects, supporting its utility as an adjunct to standard chemotherapy [10–12]. We previously demonstrated that recombinant D29 LysB exhibits direct antimycobacterial activity against drug-resistant *M. tuberculosis* isolates and enhances the activity of conventional anti-tubercular drugs *in vitro* [12]. However, the *in vivo* therapeutic potential of pulmonary LysB delivery in tuberculosis has not been evaluated. Although *in vivo* evidence remains limited, available findings are encouraging. Subcutaneous administration of LysB inhibited *M. ulcerans* growth in mice [11], whereas intrapulmonary delivery reduced *M. abscessus* burden in experimental infection models [10]. Together, these observations provide a compelling rationale for further investigation of LysB as a therapeutic candidate against TB.

Despite these promising findings, the therapeutic potential of LysB in animal models of TB remains largely unexplored, limiting translational advancement toward clinical application. Intranasal delivery represents an attractive approach for pulmonary TB because it enables high local drug concentrations in the lungs, promotes retention within alveolar macrophages, and minimizes first-pass metabolism [13, 14]. Previous studies have successfully employed intranasal delivery of vaccines, proteins, and nanoparticle-based therapeutics in TB models with favorable outcomes [15]. Intranasal lysin formulations have shown efficacy against respiratory pathogens, including *Streptococcus pneumoniae*, *Pseudomonas aeruginosa*, and *Mycobacterium abscessus* [10, 16–18]. However, despite the biological relevance of this route, the *in vivo* efficacy of intranasally delivered phage-derived endolysins against *M. tuberculosis* has not been established.

In the present study, we evaluated the therapeutic efficacy of intranasally administered recombinant D29 LysB in murine and guinea pig models of pulmonary TB. We hypothesized that localized pulmonary delivery of LysB would enhance antimycobacterial activity, improve histopathological outcomes, and beneficially modulate host immune responses without inducing adverse immunogenicity. To test this, we assessed the effects of LysB alone and in combination with standard anti-tubercular therapy (ATT) on bacterial burden, tissue pathology, cytokine responses, and antibody production. This study provides *in vivo* support for intranasal enzyme-based therapeutics as a promising adjunctive strategy for drug-resistant tuberculosis.

## Material & Method

### Bacterial strains and growth conditions

*M. tuberculosis* H37Rv was obtained from the ICMR-NJIL&OMD repository (Agra, India) and cultured in Middlebrook 7H9 medium (Difco, USA) supplemented with 10% albumin–dextrose–catalase (ADC; Difco, USA) at 37°C. Following a single passage in mice, bacterial stocks were prepared in aliquots and stored at −80°C until use. For aerosol infection experiments, frozen aliquots were thawed and subcultured to logarithmic phase in Middlebrook 7H9 broth supplemented with 10% oleic acid–albumin–dextrose–catalase (OADC; Difco, USA) and 0.05% Tween 80 (Sigma-Aldrich, USA).

### Antibiotics and LysB preparation

Isoniazid, pyrazinamide, and rifampicin were obtained from commercial sources (Sigma-Aldrich, St. Louis, MO, USA; MP Biomedicals, France). Drug solutions were prepared in distilled water and administered in final volumes of 0.2 mL for mice and 0.5 mL for guinea pigs. Fresh working solutions were prepared weekly and stored at 4°C until use. Recombinant lyophilized LysB derived from mycobacteriophage D29 was produced by heterologous expression in *Escherichia coli* BL21(DE3) (Novagen), followed by purification as previously described [12]. Before administration, lyophilized LysB was reconstituted in phosphate-buffered saline (PBS), quantified using a bicinchoninic acid (BCA) protein assay kit (Sigma-Aldrich, USA), and prepared fresh daily at the required dose for intranasal administration.

### Animals

Healthy male Hartley guinea pigs (3–3.5 months old; ∼350 g) and female BALB/c mice (8–10 weeks old, ∼23 g) were procured from the ICMR-NJIL&OMD Animal House facility (Agra, India). Animals were acclimatized for 7 days under Animal Biosafety Level 3 (ABSL-3) containment conditions prior to the initiation of experiments, with *ad libitum* access to standard pellet food and water. All animal procedures were conducted in accordance with Institutional Animal Care and Use Committee (IACUC) guidelines and were approved by the Institutional Animal Ethics Committee of ICMR-NJIL&OMD, Agra [NJIL&OMD/4IAEC/2020-01 dated 11.05.2022].

### Aerosol infections

Thirty male guinea pigs and forty-eight male BALB/c mice were housed under barrier conditions in an Animal Biosafety Level 3 (ABSL-3) facility throughout the study. Pulmonary tuberculosis infection was established by aerosol infection with frozen aliquots of *M. tuberculosis* H37Rv using an Inhalation Exposure System (Glas-Col, Terre Haute, IN, USA), as previously described [19, 20]. The exposure system was calibrated to deliver approximately 2 log^10^ CFU per animal. To verify successful inoculation, a subset of four animals was sacrificed 24 hours after infection to determine lung bacillary burden. An additional four animals were evaluated at 4 weeks post-infection to confirm the establishment of *M. tuberculosis* infection prior to treatment initiation. Animals were then randomly assigned to treatment groups. At the start of therapy, pulmonary bacterial loads had stabilized at approximately 6 log^10^ CFU/g of lung tissue in BALB/c mice and 5 log^10^ CFU in guinea pigs, indicating pulmonary tuberculosis infection (Figure 1).

**FIGURE 1.**
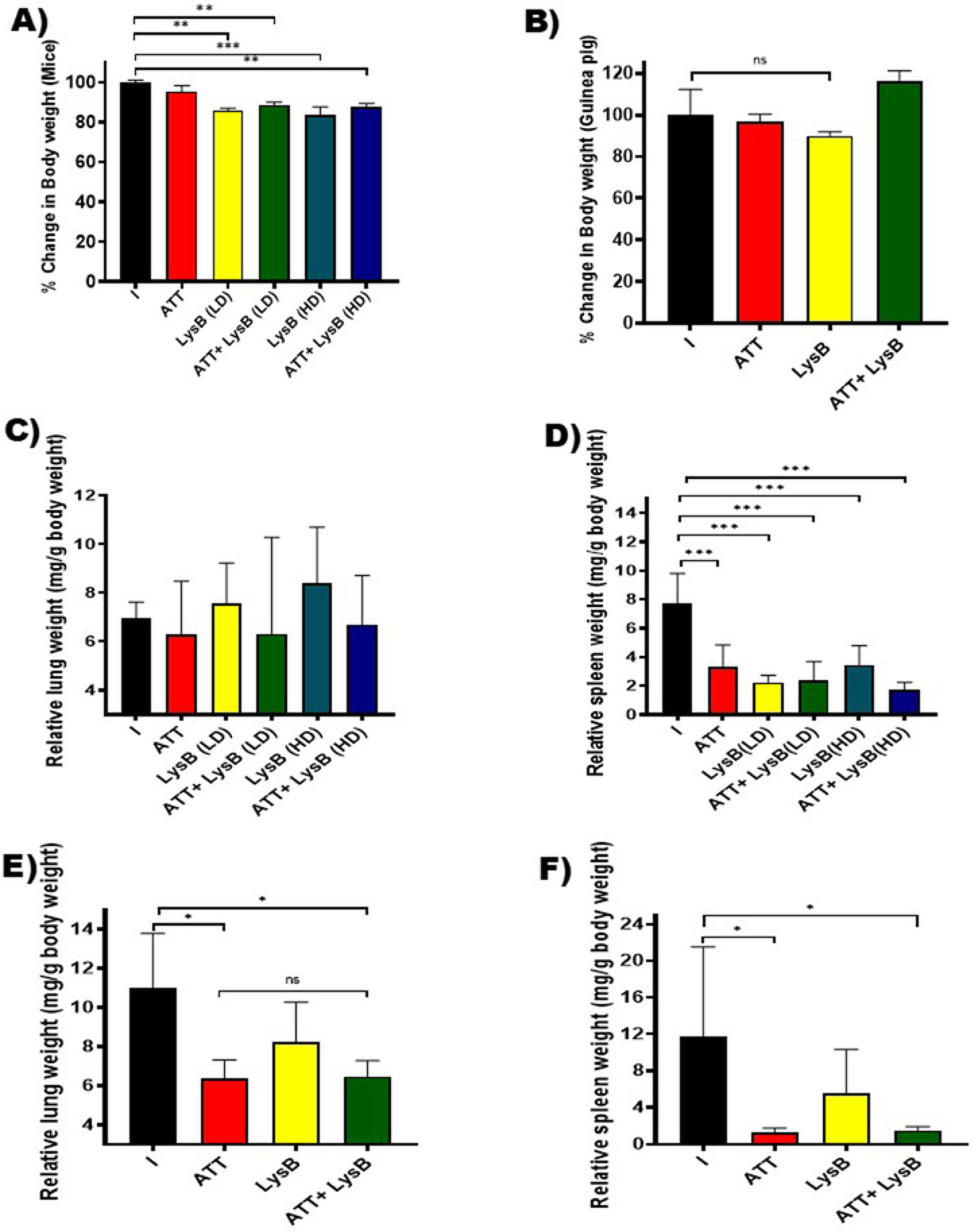
Effect of anti-tubercular therapy (ATT) and recombinant D29 LysB on body weight and normalized organ weights in *M. tuberculosis*–infected mice and guinea pigs. (A and B) Average percentage change in body weight in BALB/c mice **(A)** and guinea pigs (**B**) at the experimental endpoint relative to initial body weight following treatment with ATT, recombinant D29 LysB, or combination therapy. Mice received recombinant D29 LysB intranasally at low dose (LD; 20 µM in 50 µL) or high dose (HD; 40 µM in 50 µL), administered 5 days/week for 4 weeks. Guinea pigs received recombinant D29 LysB intranasally at 40 µM in a total volume of 100 µL on the same schedule. ATT consisted of rifampicin, isoniazid, and pyrazinamide administered orally. **(C and D)** Lung and spleen weights normalized against body weight of BALB/c mice following aerosol infection with *M. tuberculosis* H37Rv and treatment with ATT, recombinant D29 LysB, or combination therapy. **(E and F)** Lung and spleen weights normalized against the body weight of the guinea pig following treatment with ATT, recombinant D29 LysB, or combination therapy. Data are presented as mean ± SD, with n = 5 mice/group **and** n = 4 guinea pigs/group. Statistical significance was determined by one-way ANOVA followed by Dunnett’s multiple comparisons test. *, *P* < 0.05; **, *P* < 0.01; ***, P<0.001 versus untreated infected controls.

### Antibiotic treatment

Drug treatment began 28 days after aerosol infection, once chronic infection was established. Treatment continued 5 days per week for 4 weeks. Control animals received no treatment or standard short-course anti-tubercular therapy (ATT). Drugs were administered orally using a ball-tipped 20-gauge gavage cannula. Mice received rifampicin (10 mg/kg), isoniazid (25 mg/kg), and pyrazinamide (150 mg/kg) [21]. Guinea pigs received rifampicin (50 mg/kg), isoniazid (30 mg/kg), and pyrazinamide (150 mg/kg) by oral gavage, as previously described [22, 23]. These dosing regimens were selected to match drug exposures in humans receiving standard therapeutic doses. Body weights were recorded weekly during treatment. Rifampicin was given at least 1 h before the other anti-tubercular drugs in both species. Experimental groups received recombinant D29 LysB intranasally in phosphate-buffered saline (PBS). D29 LysB was administered either alone or with ATT. Mouse doses of recombinant D29 LysB were selected based on previously published *in vivo* studies evaluating pulmonary delivery of D29 LysB in experimental mycobacterial infection [10]. In mice, endotoxin-free recombinant D29 LysB was administered intranasally at concentrations of 20 or 40 µM in a total volume of 50 µL, 5 days per week for 4 weeks. Guinea pigs received 40 µM endotoxin-free D29 LysB in a total volume of 100 µL following the same treatment schedule. D29 LysB was delivered dropwise to the anterior nares using a micropipette and inhaled spontaneously by the animals.

### Study endpoints and tissue processing

The experimental design is summarized in **Table 1** and **figure 2A**. To determine the initial pulmonary bacterial implantation and bacillary burden before treatment initiation, subsets of mice and guinea pigs were sacrificed at 24 h and 4 weeks after aerosol infection, respectively. Therapeutic efficacy of standard anti-tubercular therapy (ATT; rifampicin, isoniazid, and pyrazinamide) and recombinant D29 LysB, administered alone or in combination, was evaluated after 4 weeks of treatment. At the end of the treatment period, animals were maintained for an additional 3-day drug washout period prior to sacrifice. Mice and guinea pigs were euthanized under isoflurane anesthesia according to institutional ethical guidelines. Blood samples were collected by cardiac puncture before necropsy. Lungs and spleens were aseptically removed, weighed, photographed, and processed for microbiological and histopathological analysis.

**FIGURE 2.**
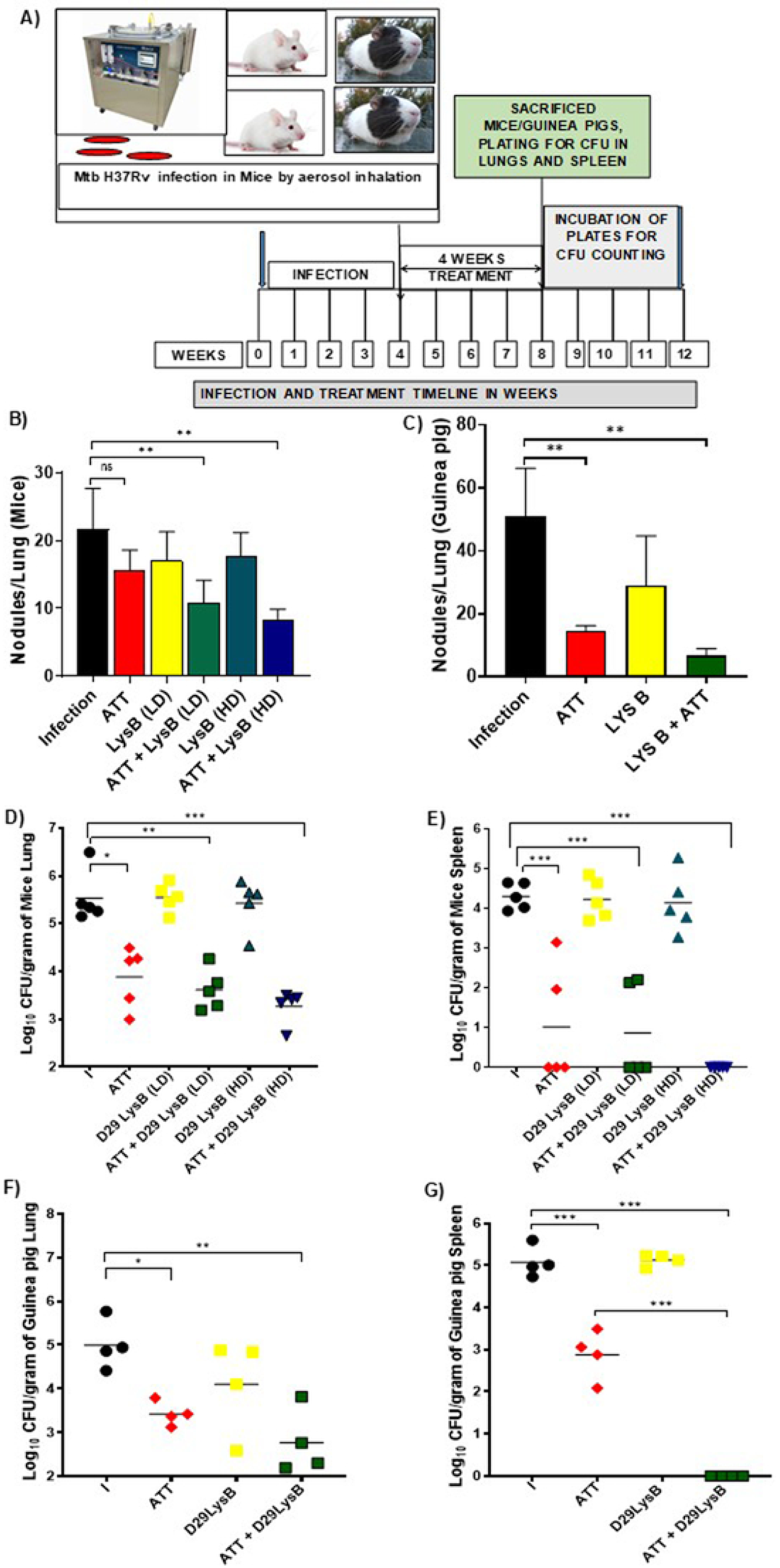
Effect of anti-tubercular therapy (ATT) and recombinant D29 LysB on pulmonary lesion burden and bacterial load in experimental tuberculosis. **(A)** Schematic illustration of experimental design **(B and C).** Morphometric quantification of granulomatous nodule numbers per lung in BALB/c mice **(B)** and guinea pigs **(C)** following aerosol infection with *M. tuberculosis* H37Rv and treatment with ATT, recombinant D29 LysB, or combination therapy. Mice received recombinant D29 LysB intranasally at low dose (LD; 20 µM in 50 µL) or high dose (HD; 40 µM in 50 µL), administered 5 days/week for 4 weeks. Guinea pigs received recombinant D29 LysB intranasally at 40 µM in a total volume of 100 µL on the same schedule. Granulomatous nodules were enumerated from representative gross lung images using Fiji (ImageJ) software. **(D and E)** Lung and spleen bacterial burden in BALB/c mice following treatment with ATT, recombinant D29 LysB, or combination therapy. **(F and G)** Lung and spleen bacterial burden in the guinea pig following treatment. Each symbol represents one animal, and horizontal bars indicate mean values. Data were analyzed after 4 weeks of treatment. Statistical significance was determined by one-way ANOVA followed by Dunnett’s multiple comparisons test. *, *P* < 0.05; **, *P* < 0.01; ***, *P* < 0.001; ****, *P* < 0.0001 versus untreated infected controls.

**Table 1.** Experimental infection and treatment schedule for murine and guinea pig models of pulmonary tuberculosis. Each experimental group consisted of six mice or five guinea pigs. Animals were aerosol infected with *M. tuberculosis* H37Rv on Day 0 and euthanized at the indicated experimental endpoints.

| Group | Experimental group | Experimental details / treatment regimen | Treatment duration | Terminal sacrifice (TS) |
| --- | --- | --- | --- | --- |
| **A** | Uninfected control | Uninfected animals receiving no treatment | — | Day 56 |
| **B** | Initial implantation control | Aerosol infection with *M. tuberculosis* H37Rv; animals sacrificed 24 h post-infection for determination of initial pulmonary bacterial burden | — | Day 1 |
| **C** | Pretreatment infection control | Aerosol infection; animals sacrificed 4 weeks post-infection to determine pulmonary bacillary burden before treatment initiation | — | Day 28 |
| **D** | Untreated infection control | Aerosol infection; no treatment administered; animals sacrificed 8 weeks post-infection to assess disease progression | — | Day 56 + 3-day washout |
| **E** | ATT-treated | Standard anti-tubercular therapy (ATT) administered orally, 5 days/week for 4 weeks. Mice: isoniazid (25 mg/kg), rifampicin (10 mg/kg), pyrazinamide (150 mg/kg). Guinea pigs: isoniazid (30 mg/kg), rifampicin (50 mg/kg), pyrazinamide (150 mg/kg). | 20 doses over 28 days | Day 56 + 3-day washout |
| **F** | LysB-treated | Recombinant D29 LysB administered intranasally, 5 days/week for 4 weeks. Mice: 20 or 40 μM in a total volume of 50 μL. Guinea pigs: 40 μM in a total volume of 100 μL. | 20 doses over 28 days | Day 56 + 3-day washout |
| **G** | Combination therapy | Standard anti-tubercular therapy combined with intranasal recombinant D29 LysB at the above doses, administered 5 days/week for 4 weeks | 20 doses over 28 days | Day 56 + 3-day washout |

**Abbreviations:** ATT, anti-tubercular therapy; TS, terminal sacrifice.

In guinea pigs, the right lung and half of the spleen were homogenized and plated on Middlebrook 7H11 agar (Becton Dickinson, Sparks, MD, USA) for colony-forming unit (CFU) enumeration, while the remaining tissue was fixed for histological examination. In mice, representative portions of lung and spleen tissues were reserved for histology, and the remaining tissue was processed for microbiological analysis. CFUs were enumerated after 21–28 days of incubation and expressed as log^10^-transformed values. Data were generated from five mice and four guinea pigs per group at each experimental endpoint.

### Gross organ morphology, morphometric analysis, and histopathology

Animal body weights and lung and spleen weights were recorded at the time of sacrifice, and organs were photographed for gross pathological assessment. To account for changes in body weight during infection and treatment, organ weights were normalized using the following formula:

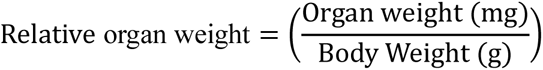

The excised lung and spleen tissues were fixed in 4% paraformaldehyde, embedded in paraffin, and sectioned at 5 μm with a rotary microtome. Sections were then stained with hematoxylin and eosin (H&E), mounted in DPX, and examined by light microscopy for pathological assessment. Histopathological evaluation focused on lung consolidation and splenic white pulp depletion as indicators of disease severity and treatment response. Pathology was scored semi-quantitatively using the following scale: (–) no detectable lesion, (+) mild pathology, (++) moderate pathology, (+++) severe pathology, and (++++) very severe pathology. All slides were evaluated by a pathologist blinded to treatment allocation. For morphometric assessment, the number of visible lung lesions was counted from representative images using Fiji (ImageJ) software.

### Quantification of cytokine levels in lung homogenates

Lung tissues from experimental groups were homogenized in phosphate-buffered saline (PBS) to a final volume of 1.5 mL using a Polytron tissue homogenizer. The homogenates were centrifuged at 10,000 × g for 10 minutes at 4°C to remove debris. The supernatants were collected and stored at −80°C until analysis. Pro-inflammatory cytokine TNF-α and anti-inflammatory cytokine IL-10 levels were measured in lung homogenate supernatants using mouse ELISA kits (R&D Systems, Minneapolis, MN, USA), following the manufacturer’s protocol with minor optimizations as previously described [24]. Absorbance was read at 450 nm with background correction at 570 nm on a BioTek Epoch 2. Samples were analyzed in triplicate. Cytokine concentrations were calculated from standard curves generated for each assay, using the absorbance values of known standards plotted against their concentrations to interpolate sample concentrations. Results were expressed as pg/mL.

### Enzyme-linked immunosorbent assay (ELISA) for the detection of D29 LysB-specific antibodies

Recombinant D29 LysB was coated onto 96-well ELISA plates (NEST, India) at 5 ng/well in carbonate coating buffer (Na_2_CO_3_, pH 8.0) and incubated overnight at 4°C. After four washes with PBST (PBS containing 0.05% Tween-20), the plates were blocked with PBSM (1% bovine serum albumin in PBS) for 1 h at room temperature. Plates were then washed three times with PBST, and mouse serum samples, diluted 1:100 in PBSM, were added (100 µL/well) and incubated for 2 h at room temperature. Following incubation, plates were washed five times with PBST, then incubated for 1 h at room temperature with 100 µL of HRP-conjugated secondary antibodies. For detection of LysB-specific antibody responses, HRP-conjugated rat anti-mouse IgG and goat anti-mouse IgE antibodies (Invitrogen) were used. After a final wash, tetramethylbenzidine (TMB) substrate was added to develop color, and absorbance was measured at 450 nm with 570 nm background correction using a BioTek Synergy 2.

### Statistical analysis

Bacterial burden data, expressed as colony-forming units (CFU), were log-transformed using log10(x + 1) prior to analysis. Differences in bacterial load and cytokine and antibody response data among experimental groups were analyzed by one-way analysis of variance (ANOVA) followed by Dunnett’s multiple comparisons test, with each treatment group compared against the untreated infected control. Statistical analyses were performed using GraphPad Prism (GraphPad Software, San Diego, CA, USA). Data are presented as mean ± standard deviation (SD), and differences were considered statistically significant at P < 0.05.

## Results

### Treatment-Associated Morbidity and Body Weight Changes in Mice and Guinea Pigs

Throughout the experiments, no deaths occurred in either the experimental or control group of mice or guinea pigs. However, mice receiving the treatment experienced weight reduction without noticeable changes in their overall behavior. The ATT-treated mice group showed a weight decrease of approximately 1.7%, which was more pronounced in the groups treated with LysB (≥10%) **(Figure 1A)**. Mice receiving ATT in combination with LysB exhibited approximately 9% body weight loss during the treatment period. Comparable weight reduction (<10%) has previously been reported following repeated D29 LysB administration in murine infection models, suggesting that this observation may be associated with repeated enzyme administration rather than overt treatment-related toxicity [10]. Weight loss in ATT-treated mice may reflect treatment-associated handling stress or daily drug administration in 200 µL volumes by oral route, potentially limiting their food consumption [25].

In contrast to mice, guinea pigs treated with LysB alone exhibited progressive weight gain during treatment. Animals receiving combination therapy with ATT and LysB also showed increased body weight and did not exhibit treatment-associated weight loss at any point during the experiment. At the experimental endpoint, guinea pigs treated with ATT plus LysB demonstrated significantly greater body weight gain (≥16%) compared with untreated infected controls and ATT-treated animals **(Figure 1B)**. No TB-associated morbidity or treatment-related adverse effects were observed in guinea pigs during the study period..

### Organ weights, gross pathology, and histology during treatment

Relative lung and spleen weights (mg/g body weight) were measured at the experimental endpoint to evaluate disease-associated organ pathology **(Figure 1C–F)**. In mice, relative lung weights showed only minor differences among treatment groups **(Figure 1C)**. Untreated, infected mice had increased relative spleen weights compared with treated groups **(Figure 1D)**. ATT treatment and combination therapy with ATT and D29 LysB reduced spleen weights significantly (p<0.001), with combination therapy showing the greatest reduction. LysB monotherapy led to only partial improvement.

In guinea pigs, untreated infected animals exhibited elevated relative lung and spleen weights, consistent with progressive pulmonary and systemic disease **(Figure 1E–F)**. ATT treatment significantly reduced relative lung and spleen weights compared with untreated controls. Combination therapy with ATT and D29 LysB also reduced relative organ weights and demonstrated a trend toward greater reduction in lung and spleen weight compared with ATT alone. LysB monotherapy produced comparatively limited effects.

### Morphometric Assessment of Pulmonary Granulomatous Lesions

Gross examination of lungs from untreated *M. tuberculosis*–infected mice and guinea pigs showed extensive granulomatous involvement with numerous visible nodules on the lung surface **(Figure 2B and C)**. Quantitative morphometric analysis indicated fewer pulmonary lesions after treatment with recombinant D29 LysB. In BALB/c mice, this decrease was most pronounced in animals given anti-tubercular therapy (ATT) combined with D29 LysB, compared to those receiving ATT or LysB alone. The most significant reduction appeared in the group ATT plus high-dose LysB **(Figure 2B)**. A similar pattern occurred in guinea pigs, as combination treatment greatly reduced visible lung lesions versus untreated controls and monotherapy groups **(Figure 2C).**

### Bactericidal activity of D29LysB alone and in combination with ATT against chronic tuberculosis infection in mice

On the day following aerosol infection, 1.96 ± 0.48 log^10^ CFU were recovered from mouse lungs. The lung bacillary burden increased to 6.4 ± 0.14 log^10^ CFU by the day of treatment initiation. In untreated mice, host immune responses contained bacillary growth, resulting in a stable lung burden of 5.53 ± 0.24 log^10^ CFU at the end of the experiment **(Figure. 2D)**. Mice treated with ATT (RHZ), administered five days per week for four weeks, exhibited a 1.75 log^10^ reduction in CFU at the end of treatment, with lung CFU decreasing to 3.8 ± 0.34 log^10^ CFU after 28 days. D29LysB at 20 µM and 40 µM did not demonstrate significant bactericidal activity by day 28, as lung CFU counts in these groups were 5.54 ± 0.13 log^10^ and 5.42 ± 0.23 log^10^, respectively. By day 28, the combination therapy of ATT and D29LysB at 40 µM exhibited significantly greater activity than ATT alone at human equivalent doses, reducing the bacterial burden in mouse lungs by an additional 0.6 log^10^ compared to the ATT-treated group (3.2 ± 0.3 log^10^ CFU). Combination therapy with ATT and D29LysB at 20 µM had only a slight effect on mycobacterial survival compared to ATT treatment alone, reducing lung CFU to 3.6 ± 0.21 log^10^ CFU (**Figure 2D**). In the spleen, LysB at 20 µM or 40 µM did not reduce mycobacterial growth compared to the untreated group. ATT or ATT combined with 20 µM LysB reduced mycobacterial growth by 3 log^10^ CFU relative to untreated mice. Notably, the combination of 40 µM LysB with ATT cleared mycobacterial infection in all mice (5/5). In contrast, ATT alone cleared infection in only 2 out of 5 mice (**Figure 2E**).

### Evaluation of the bactericidal activity of D29 LysB alone and in combination with ATT in a guinea pig model of chronic tuberculosis infection

Guinea pigs were infected via the aerosol route, resulting in a day-1 lung implantation of 1.91 ± 0.08 log^10^ CFUs. Drug treatments began four weeks post-infection. At this point, the bacterial lung burden reached 5.3 ± 0.11 log^10^ CFUs. Treatment with standard anti-tubercular therapy (ATT) for 4 weeks reduced lung bacterial burden by approximately 1.3 log^10^ CFU compared with untreated controls (**Figure 2F**). In contrast, D29 LysB monotherapy produced only a modest reduction in pulmonary bacillary load, decreasing lung CFU counts by ∼0.7 log^10^ relative to untreated animals. Adjunctive administration of D29 LysB together with ATT resulted in the greatest reduction in pulmonary bacterial burden, producing an overall ∼2 log^10^ decrease compared with untreated controls and an additional ∼0.7 log^10^ reduction relative to ATT alone (**Figure 2G**).

A similar trend was observed in the spleen (Figure 2F). ATT treatment significantly reduced splenic bacterial burden to 2.9 ± 0.42 log^10^ CFU compared with untreated animals (P ≤ 0.0001). D29 LysB alone had a limited effect on splenic bacterial counts. Notably, no culturable bacilli were recovered from the spleens of guinea pigs receiving combination therapy with ATT and D29 LysB, indicating complete clearance of detectable extrapulmonary infection at the experimental endpoint. (**Figure 2G**).

### Histopathological evaluation of lung and spleen tissues

Histopathological examination of lung and spleen tissues from *M. tuberculosis*–infected mice and guinea pigs demonstrated marked treatment-associated improvement following administration of ATT and recombinant D29 LysB **(Figures 3 and 4**; **Table 2).** Untreated infected animals showed severe pathological alterations in both organs, whereas treatment with ATT and recombinant D29 LysB reduced the severity of tissue damage to varying degrees. In mice, lung sections from untreated animals showed severe multifocal fibrinous degeneration, mononuclear cell infiltration, and caseous necrosis (score: +++), together with diffuse fibrin accumulation (score: +++). ATT treatment reduced the severity of these lesions to moderate levels (++). LysB monotherapy also reduced pulmonary pathology, with less severe inflammatory infiltration and tissue degeneration compared with untreated controls. The greatest improvement was observed in animals receiving ATT together with LysB, particularly at the higher dose, where most pulmonary lesions were reduced to mild severity (+). Marked splenic pathology was also evident in untreated mice, including depletion of red and white pulp, vacuolar degeneration, and caseous changes. These abnormalities were less pronounced in ATT- and LysB-treated groups. Combination treatment with ATT and LysB resulted in minimal or undetectable splenic pathological changes, with histological appearance approaching that of normal tissue.

**FIGURE 3.**
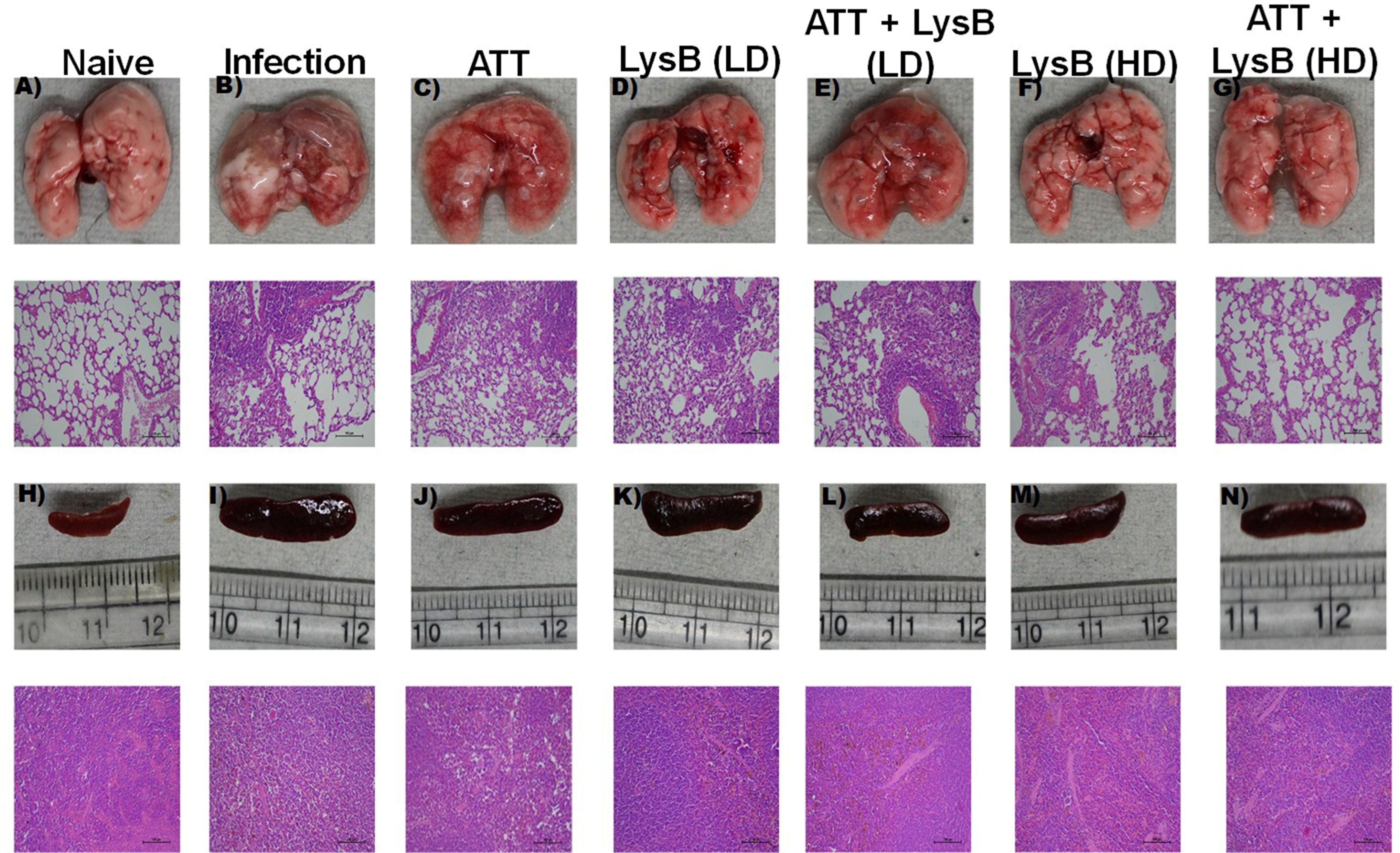
Gross morphological and histopathological analysis of lungs and spleens from *M. tuberculosis*–infected BALB/c mice following treatment with ATT and intranasal recombinant D29 LysB. Low-dose (LD) D29 LysB was administered intranasally at 20 µM in 50 µL, whereas high-dose (HD) D29 LysB was administered at 40 µM in 50 µL, 5 days/week for 4 weeks. ATT consisted of rifampicin, isoniazid, and pyrazinamide administered orally. **Top row (A–G):** Representative gross morphology of lungs collected from BALB/c mice following aerosol infection with *M. tuberculosis* H37Rv and 4 weeks of treatment with standard anti-tubercular therapy (ATT), recombinant D29 LysB (20 or 40 µM, intranasal), or combination therapy. **Second row:** Representative hematoxylin and eosin (H&E)-stained lung sections corresponding to the groups shown in panels A–G. **Third row (H–N):** Representative gross morphology of spleens collected from corresponding experimental groups. **Bottom row:** Representative H&E-stained spleen sections corresponding to the groups shown in panels H–N. **(A and H)** Naïve control. (B and I) *M. tuberculosis*–infected, untreated control. **(C and J)** ATT-treated group. **(D and K)** low-dose (LD) D29 LysB-treated group. **(E and L)** high-dose (HD) D29 LysB-treated group. **(F and M)** ATT + LD D29 LysB-treated group. **(G and N)** ATT + HD D29 LysB-treated group.

**FIGURE 4.**
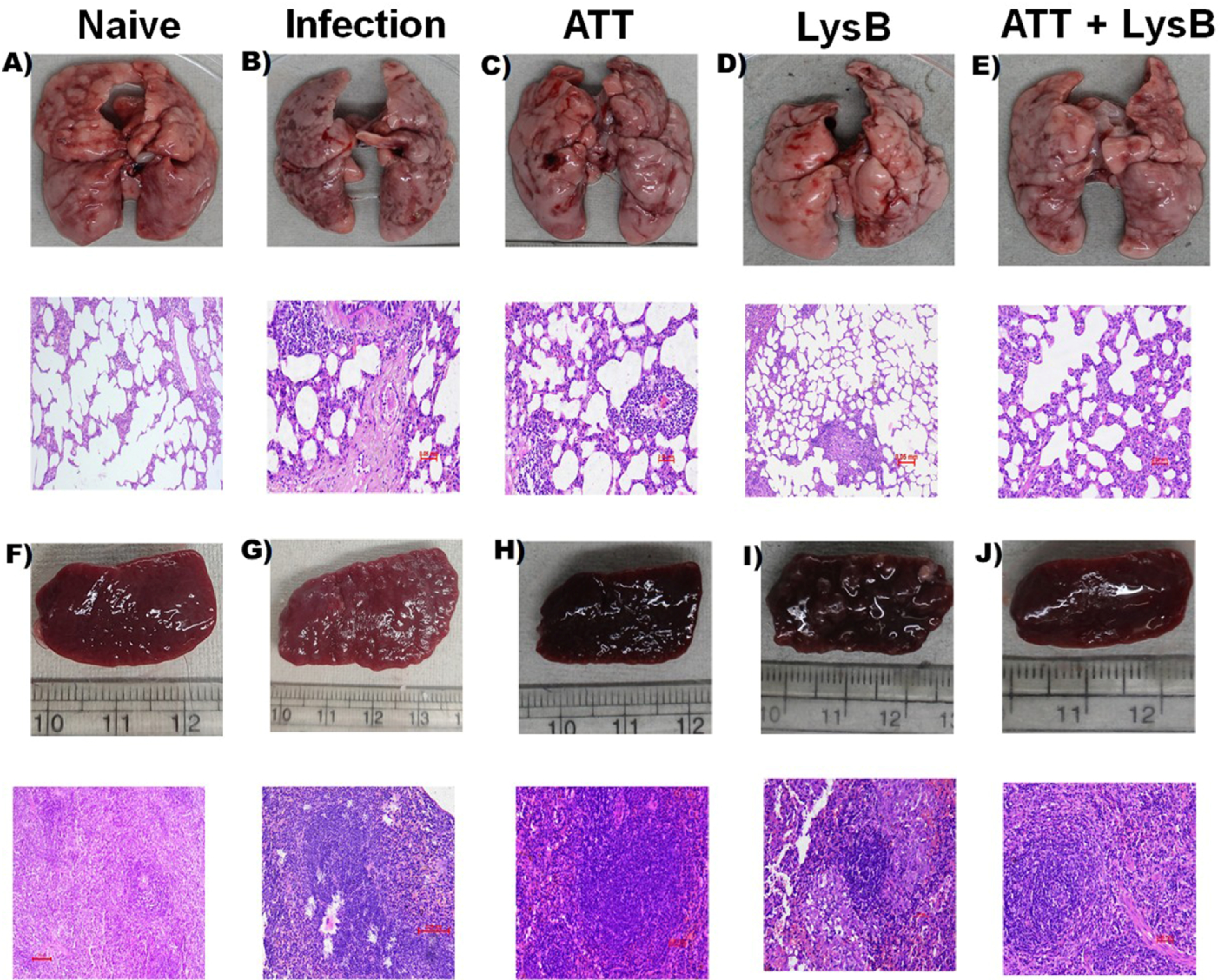
Gross morphological and histopathological analysis of lungs and spleens from *M. tuberculosis*–infected guinea pigs following treatment with ATT and intranasal D29 LysB. Lung, upper panels: Representative gross morphology of lungs collected from guinea pigs after aerosol infection with *M. tuberculosis* H37Rv and following 4 weeks of treatment with standard anti-tubercular therapy (ATT), recombinant D29 LysB (40 µM, intranasal), or combination therapy. **Lung, lower panels:** Representative hematoxylin and eosin (H&E)-stained lung sections from corresponding treatment groups. **Spleen, upper panels:** Representative gross morphology of spleens collected from corresponding experimental groups. **Spleen, lower panels:** Representative H&E-stained spleen sections from corresponding treatment groups. **(a)** Naïve control. **(b)** *M. tuberculosis*–infected, untreated control. **(c)** ATT-treated group. **(d)** D29 LysB (40 µM, intranasal)-treated group. **(e)** ATT + D29 LysB (40 µM /kg, intranasal)-treated group.

**Table 2.** Histopathological scoring of lung and spleen lesions in murine and guinea pig models of pulmonary tuberculosis following treatment with ATT and recombinant D29 LysB.

| Animal model | Tissue | Histopathological observation | Control | ATT | LysB (20 μM) | ATT + LysB | LysB (40 μM) | ATT + LysB |
| --- | --- | --- | --- | --- | --- | --- | --- | --- |

|  |  |  |  |  |  | (20<br>μM) |  | (40<br>μM) |
| --- | --- | --- | --- | --- | --- | --- | --- | --- |
| **Mouse** | Lung | Multifocal fibrinous degeneration | +++ | ++ | +++ | ++ | ++ | + |
|  |  | Multifocal mononuclear cell infiltration | +++ | ++ | +++ | ++ | ++ | + |
|  |  | Caseous necrosis | +++ | ++ | +++ | ++ | ++ | + |
|  |  | Diffuse fibrin accumulation | ++ | ++ | +++ | ++ | ++ | + |
|  | Spleen | Decrease in red pulp with giant cells | +++ | — | + | — | + | — |
|  |  | Vacuolar degeneration | +++ | — | + | — | + | — |
|  |  | Caseous necrosis | ++ | — | + | — | + | — |
|  |  | Decrease in white pulp | +++ | — | + | — | + | — |
|  |  | Decrease in hemosiderosis | ++ | — | + | — | + | — |
| **Guinea pig** | Lung | Multifocal fibrinous degeneration | +++ | ++ | ND | ND | +++ | + |
|  |  | Multifocal mononuclear cell infiltration | +++ | ++ | ND | ND | +++ | + |
|  |  | Diffuse fibrin accumulation | +++ | ++ | ND | ND | +++ | + |
|  |  | Primary granuloma | +++ | ++ | ND | ND | +++ | + |
|  |  | Secondary granuloma | +++ | ++ | ND | ND | +++ | + |
|  |  | Granuloma disruption | +++ | ++ | ND | ND | +++ | + |
|  | Spleen | Decrease in red pulp with giant cells | ++++ | ++ | ND | ND | ++ | + |
|  |  | Vacuolar degeneration | ++++ | ++ | ND | ND | ++ | + |
|  |  | Decrease in white pulp | ++++ | ++ | ND | ND | ++ | + |
|  |  | Decrease in hemosiderosis | ++++ | ++ | ND | ND | ++ | + |

**Scoring criteria:** –, absent; +, mild; ++, moderate; +++, severe; +++++, very severe
histopathological changes.
**Abbreviations:** ATT, anti-tubercular therapy; ND; not done

A similar pattern was observed in guinea pigs. Untreated infected animals exhibited severe pulmonary lesions characterized by fibrinous degeneration, mononuclear cell infiltration, granulomatous pathology, and disruption of lung architecture (score: +++). ATT treatment reduced lesion severity, while LysB monotherapy produced partial histological improvement. Combination treatment with ATT and LysB reduced most pulmonary lesions to mild severity (+).

In the spleen, untreated guinea pigs showed marked depletion of red and white pulp with vacuolar degeneration and giant cell infiltration (score: ++++). These changes were moderately reduced following ATT or LysB monotherapy (++), whereas combination treatment produced only mild residual splenic pathology (+), indicating substantial histological recovery.

### Cytokine Responses to D29LysB in Lung Homogenates

Lung homogenates from animals subjected to different treatment regimens were analyzed using ELISA to quantify pro-inflammatory and anti-inflammatory cytokines. Tumor necrosis factor-alpha (TNF-α), a key pro-inflammatory cytokine essential for macrophage activation and granuloma maintenance in tuberculosis, exhibited variable modulation across experimental groups. In all treated animals, TNF-α expression was suppressed to differing degrees compared to untreated controls. This pattern aligns with the immune evasion strategy of *M. tuberculosis*, which reduces pro-inflammatory responses during chronic infection to enhance its survival. Treatment with standard anti-tubercular therapy (ATT) and a low dose of LysB (20 µM) caused a modest, non-significant increase in TNF-α compared to infected, untreated animals. Mice treated with LysB at 40 µM alone showed a similar trend. Notably, the combination of ATT with LysB led to a greater increase in TNF-α than either treatment alone (*P* < 0.05) **(Figure 5A)**. Simultaneously, levels of the Type 2 cytokine interleukin-10 (IL-10) were modestly elevated across all treatment groups, although these changes did not reach statistical significance. The highest IL-10 concentrations were observed in the group receiving LysB (20 µM) in combination with ATT **(Figure 5B)**.

**FIGURE 5.**
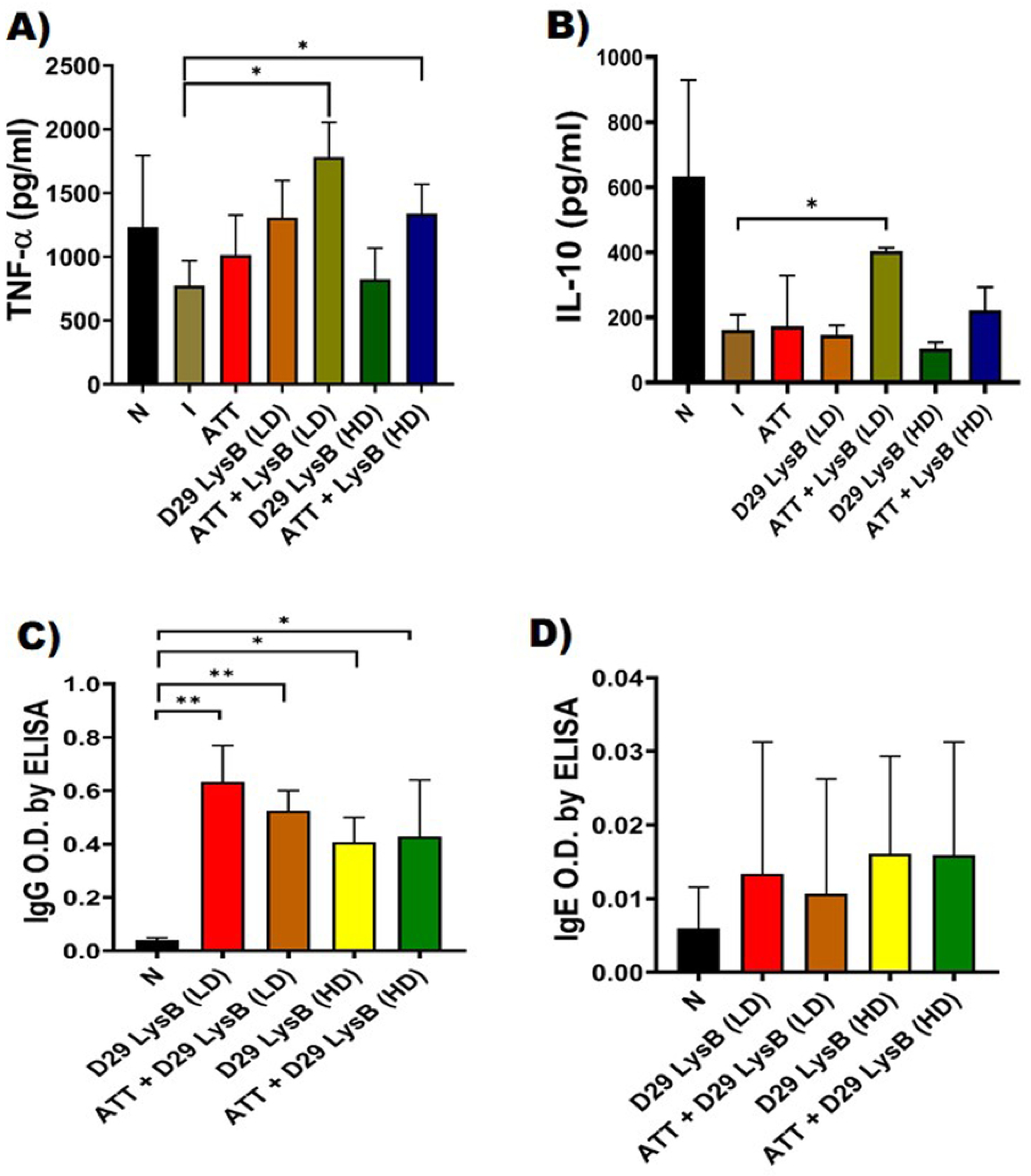
Effect of recombinant D29 LysB treatment on cytokine responses and humoral immune responses in *M. tuberculosis*–infected mice. (A) TNF-α levels in lung homogenates from BALB/c mice following treatment with standard anti-tubercular therapy (ATT), recombinant D29 LysB, or combination therapy. **(B)** IL-10 levels in lung homogenates from corresponding treatment groups. **(C)** LysB-specific IgG responses measured by ELISA following repeated intranasal administration of recombinant D29 LysB alone or in combination with ATT. **(D)** LysB-specific IgE responses measured by ELISA in corresponding treatment groups. Data are presented as mean ± SD, with n = 5 mice/group. Statistical significance was determined by one-way ANOVA followed by Dunnett’s multiple comparisons test. *, *P* < 0.05; **, *P* < 0.01 versus untreated infected controls. ***Abbreviations:*** ATT, anti-tubercular therapy; low dose (LD; 20 µM in 50 µL, high dose (HD; 40 µM in 50 µL)

### Humoral Immune Response Following Repeated LysB Administration

LysB-specific humoral responses were measured by ELISA to detail the immune response to repeated intranasal LysB administration. **Figure 5C** shows that recombinant D29 LysB treatment led to a significant rise in antigen-specific IgG levels compared with naïve controls. This result confirms immune recognition of the administered protein. Both low-dose and high-dose LysB monotherapy groups showed similar IgG responses. Increasing the dose did not substantially enhance antibody production. Co-administration of anti-tubercular therapy (ATT) with LysB produced lower IgG levels than LysB monotherapy. No significant difference was detected between the ATT plus LysB low-dose and high-dose groups. This suggests that ATT modulates humoral responses regardless of the LysB dose.

LysB-specific IgE levels stayed low for all treatment groups, with no significant differences compared to naïve controls (**Figure 5D**). The absence of elevated IgE responses suggests that repeated intranasal LysB exposure did not cause IgE-mediated hypersensitivity. This supports a favorable immunological safety profile.

## Discussion

The emergence of multidrug-resistant and extensively drug-resistant tuberculosis has increased the need for therapeutic approaches that improve the activity of existing chemotherapy through mechanisms distinct from conventional antibiotics. In the present study, intranasal administration of recombinant mycobacteriophage-derived D29 LysB reduced pulmonary and splenic bacillary burden and improved tissue pathology in both murine and guinea pig models of tuberculosis. Although LysB monotherapy showed only modest bactericidal activity, adjunctive administration with standard anti-tubercular therapy (ATT) consistently enhanced bacterial clearance, including complete elimination of detectable splenic bacilli in several animals. These microbiological findings were accompanied by improved preservation of lung and splenic architecture, supporting the therapeutic benefit of combining LysB with conventional chemotherapy.

The present findings extend previous *in vitro* studies, including our group, demonstrating antimycobacterial activity of D29 LysB against *M. abscessus*, *M. ulcerans*, and drug-resistant *M. tuberculosis* isolates [8, 10–12]. LysB functions as a mycolylarabinogalactan esterase that hydrolyzes the linkage between mycolic acids and arabinogalactan within the mycobacterial cell envelope [8]. Disruption of this highly lipid-rich permeability barrier may enhance penetration or activity of first-line anti-tubercular drugs within infected tissues. Intranasal delivery additionally permits direct deposition of the enzyme within the respiratory tract while avoiding first-pass hepatic metabolism. Previous studies using inhaled phage preparations against *M. abscessus* and *Pseudomonas aeruginosa* infections have similarly reported improved local efficacy following pulmonary delivery [26].

In both murine and guinea pig models, adjunctive LysB treatment produced an additional reduction of approximately 0.6-0.7 log^10^ CFU beyond that achieved with ATT alone, corresponding to roughly a 4 to 5-fold decrease in bacterial burden. This effect is directionally consistent with the rapid antimycobacterial activity reported by Ojha et al., who observed approximately a one log^10^ reduction in *M. abscessus* viability after one week of pulmonary LysB treatment, although differences in pathogen, delivery route, and experimental model limit direct quantitative comparison [10]. The relatively modest magnitude of bactericidal activity observed in our study may partly reflect the limitations of intranasal protein delivery in small-animal tuberculosis models, including variable pulmonary deposition, mucociliary clearance, and incomplete penetration into infected lung regions, particularly granulomatous lesions in guinea pigs [27, 28]. These delivery constraints may have reduced the amount of active enzyme reaching relevant tissue compartments, thereby limiting apparent efficacy. Optimization of pulmonary delivery systems, such as aerosolized, liposomal, or nanoparticle-based formulations, may improve local retention and enhance therapeutic activity in future studies. Notably, the observed reduction in bacillary burden across both species is biologically meaningful because murine tuberculosis typically features more diffuse inflammatory lesions, whereas guinea pigs develop necrotizing granulomas that more closely resemble human pulmonary disease [29]. Although the magnitude of effect was modest, its reproducibility across two distinct animal models supports the translational relevance of adjunctive LysB therapy.

The reduction in splenic bacterial burden following intranasal LysB administration suggests that the therapeutic effect was not restricted solely to the pulmonary compartment. Improved pulmonary bacterial control likely reduced subsequent hematogenous dissemination of *M. tuberculosis* to extrapulmonary organs, although limited systemic exposure following intranasal administration cannot be excluded. Proteins delivered through the respiratory mucosa may undergo absorption through vascular and lymphatic pathways while retaining biological activity [30, 31]. Given the relatively small molecular size of LysB (∼29.3 kDa), transient systemic distribution after pulmonary delivery remains plausible and may contribute to the reduction in splenic bacterial burden observed in the present study. Additional pharmacokinetic and biodistribution studies will be necessary to define pulmonary retention, systemic transport, and *in vivo* enzymatic stability following intranasal administration.

Histopathological improvement in LysB-treated animals further supported the microbiological findings. Combination treatment with ATT and LysB was associated with reduced granulomatous involvement, decreased lung consolidation, and preservation of splenic architecture compared with untreated controls. These changes suggest that improved bacterial clearance occurred without evidence of excessive inflammatory tissue injury. Consistent with this interpretation, cytokine profiling demonstrated increased TNF-α together with moderate elevation of IL-10 in LysB-treated animals, particularly in combination with ATT. Similar immunomodulatory effects have previously been reported for phage-derived lysins and other cell wall-active antimicrobials in experimental infection models [11, 32]. One possible explanation is that partial degradation of the mycobacterial envelope by LysB increases exposure of immunostimulatory bacterial components within infected tissues, thereby influencing local innate immune responses.

Repeated intranasal administration of LysB was well tolerated and did not induce detectable IgE-mediated hypersensitivity. LysB-treated animals developed measurable antigen-specific IgG responses; however, antibody formation was not associated with loss of therapeutic activity. Previous studies have shown that antibodies generated against lysins often have limited effects on bacteriolytic function because of the strong affinity of lysins for their bacterial substrates [33]. Interestingly, animals receiving ATT together with LysB showed comparatively lower LysB-specific IgG responses than animals treated with LysB alone. Rifampicin-containing regimens have previously been associated with altered corticosteroid metabolism and immune responsiveness [34, 35], which may have contributed to reduced antibody production during combination therapy. Because LysB was delivered through the respiratory mucosa, additional studies examining pulmonary antigen handling, mucosal immune responses, and repeated-dose immunogenicity will be important for further translational development of inhaled LysB therapy.

The present study has several limitations. Although therapeutic efficacy was demonstrated in both murine and guinea pig models, the relatively short treatment duration and limited group sizes restrict assessment of long-term relapse prevention and chronic safety following repeated administration. In addition, the pharmacokinetic profile and tissue biodistribution of LysB were not examined directly. The study also utilized a single laboratory strain of *M. tuberculosis* and therefore may not fully reflect the phenotypic diversity of clinically relevant multidrug-resistant and extensively drug-resistant isolates. Moreover, LysB was evaluated only in its native soluble form. While the direct antimycobacterial activity of D29 LysB has previously been established in vitro [12], future studies incorporating inhalable formulation approaches, including liposomal or nanoparticle-based delivery systems, together with detailed pharmacokinetic and biodistribution analyses, may further improve pulmonary retention and therapeutic efficacy.

## Conclusion

Intranasal administration of recombinant mycobacteriophage-derived D29 LysB was well tolerated and enhanced the activity of standard anti-tubercular therapy in murine and guinea pig models of pulmonary tuberculosis. Although LysB monotherapy showed limited bactericidal activity, adjunctive treatment consistently improved mycobacterial clearance and reduced tissue pathology across both animal models. These findings support further evaluation of inhaled phage-derived endolysins as adjunctive therapeutics for tuberculosis, including drug-resistant disease. Future studies examining pulmonary pharmacokinetics, formulation optimization, and activity against clinically relevant multidrug-resistant isolates will be important for translational development of LysB-based therapy.

## Author Contributions

**Conceptualization and funding acquisition:** AKS, VJ and AM. **Methodology:** SKR, RS, RG, AKS, AM and VJ **Investigation:** SKR, RS, RG, PN, and AKS. **Data analysis and interpretation:** SKR, RS, AKS, and VJ. **Writing—original draft preparation:** SKR, AKS, AKS, AM, and VJ. **Writing—review and editing:** all authors. **Supervision:** AM, AKS and VJ.

## Acknowledgments

This work was supported by a research grant from the Indian Council of Medical Research, New Delhi (Grant No. 5/8/5/38/2019/ECD-I), awarded to Amit Kumar Singh, Amit Misra, and Vikas Jain. We are grateful to the staff of the ABSL-3 facility at ICMR-NJIL&OMD, Agra, particularly Nilesh Pal and Amit Kumar, for technical assistance during the animal experiments.

